# Midbrain Glutamatergic Neurons Modulate the Acoustic Startle Reflex and Prepulse Inhibition in Mice

**DOI:** 10.1101/2025.03.28.646009

**Authors:** Luis Enrique Martinetti, Erika Correll, Adolfo Ernesto Cuadra, Gina Castellano, Karine Fénelon

## Abstract

Prepulse inhibition (PPI) of the auditory startle reflex task is a widely recognized operational measure of sensorimotor gating. PPI deficits are a hallmark feature of schizophrenia, often associated with attentional and cognitive impairments. Despite its extensive use in preclinical research for screening antipsychotic drugs, the precise cellular and circuit mechanisms underlying PPI remain unclear, even under physiological conditions. Recent evidence suggests that non-cholinergic inputs from the pedunculopontine tegmental nucleus (PPTg) to the caudal pontine reticular nucleus (PnC) mediate PPI. In this study, we investigated the contribution of PPTg glutamatergic neurons to acoustic startle and PPI. Tract-tracing, immuno-histochemical analyses, and *in vitro* whole-cell recordings in wild-type mice confirmed that PPTg glutamatergic neurons innervate the PnC. Optogenetic inhibition of PPTg-PnC glutamatergic synapses *in vivo* resulted in increased PPI across various interstimulus intervals. Notably, while optogenetic activation of this pathway had no additional effect on startle and PPI, activation of this connection alone before startle stimulation reduced startle at short interstimulus intervals and increased startle at longer intervals. Furthermore, although PPTg glutamatergic inputs target PnC glycinergic neurons, our *in vitro* whole-cell recordings combined with optogenetic stimulation at PPTg-PnC synapses revealed that PPTg glutamatergic inputs activate PnC glutamatergic giant neurons. Our findings identify a feed-forward excitatory mechanism within the brainstem startle circuit, whereby PPTg glutamatergic inputs modulate PnC neuronal activity. These results provide new insights into the clinically relevant theoretical construct of PPI, which is disrupted in various neuropsychiatric and neurological disorders.

**Significance Statement:** The pedunculopontine tegmental nucleus (PPTg) contributes to prepulse inhibition (PPI) of startle, a translational measure of sensorimotor gating that is impaired in neuropsychiatric disorders, including schizophrenia. While PPTg cholinergic neurons have been shown not to contribute to PPI, the mechanisms by which non-cholinergic PPTg neurons modulate PPI remain incompletely understood. Here, combining tract-tracing, optogenetics, *in vitro* electrophysiology, and *in vivo* startle testing in mice, we show that PPTg glutamatergic neurons that innervate startle-mediating neurons of the caudal pontine reticular nucleus (PnC) play a critical role in PPI. Their activation modulates startle responses in a time-dependent manner, either decreasing or increasing startle magnitude. These findings refine our understanding of sensorimotor gating neural mechanisms and may inform therapeutic strategies.

## Introduction

Prepulse inhibition (PPI) of startle responses is the gold standard operational measure of sensorimotor gating, a pre-attentive mechanism by which sensory events inhibit motor outputs. (Swerdlow et al., 2000; Braff et al., 2001). PPI occurs when the presentation of a non-startling acoustic, visual or tactile stimulus prior to a startling stimulus reduces the subsequent startle response (Hoffman and Fleshler, 1963). PPI deficits are associated with and predictive of symptoms including attention impairments, psychosis and motor/speech dysfunctions. These deficits occur in several neurological disorders: schizophrenia (Mena et al., 2016; Ding et al., 2023), obsessive compulsive disorder (Swerdlow and Geyer, 1993; Kohl et al., 2015), Tourette syndrome (Castellanos et al., 1996; Buse et al., 2016), and post-traumatic stress disorder (Grillon et al., 1996). PPI exhibits high translational value because it can be elicited in human populations (Takahashi and Kamio, 2018) and other species (Koch, 1999; Burgess and Granato, 2007; Aguilar et al., 2018) and can predict psychotic symptom severity (Graham, 1975; Koch, 1999; Burgess and Granato, 2007). While PPI and the reversal of PPI deficits by antipsychotic drugs have been investigated in animal models of disease (Swerdlow and Geyer, 1993; Sinclair et al., 2017; Zheng et al., 2023), the circuit-level mechanisms underlying PPI remain unknown. Consequently, common antipsychotics show inconsistent effects on PPI in affected individuals, highlighting a crucial knowledge gap. Therefore, it is essential to identify the cellular mechanisms underlying PPI to understand the impact of disease and develop targeted therapeutic interventions.

In the mammalian acoustic brainstem startle circuit, auditory information activates cochlear root neurons which mostly activate contralateral startle-mediating giant glutamatergic neurons located in the caudal pontine reticular nucleus (PnC). Interneurons expressing the glycine transporter type 2 (GlyT2; Zeilhofer et al., 2005) are intermingled between these PnC giant neurons which directly innervate cervical and spinal motoneurons and therefore represent the sensorimotor interface of the startle pathway (**Fig. 1**; Lingenhohl and Friauf, 1992; Yeomans and Frankland, 1996).

**Figure 1.**
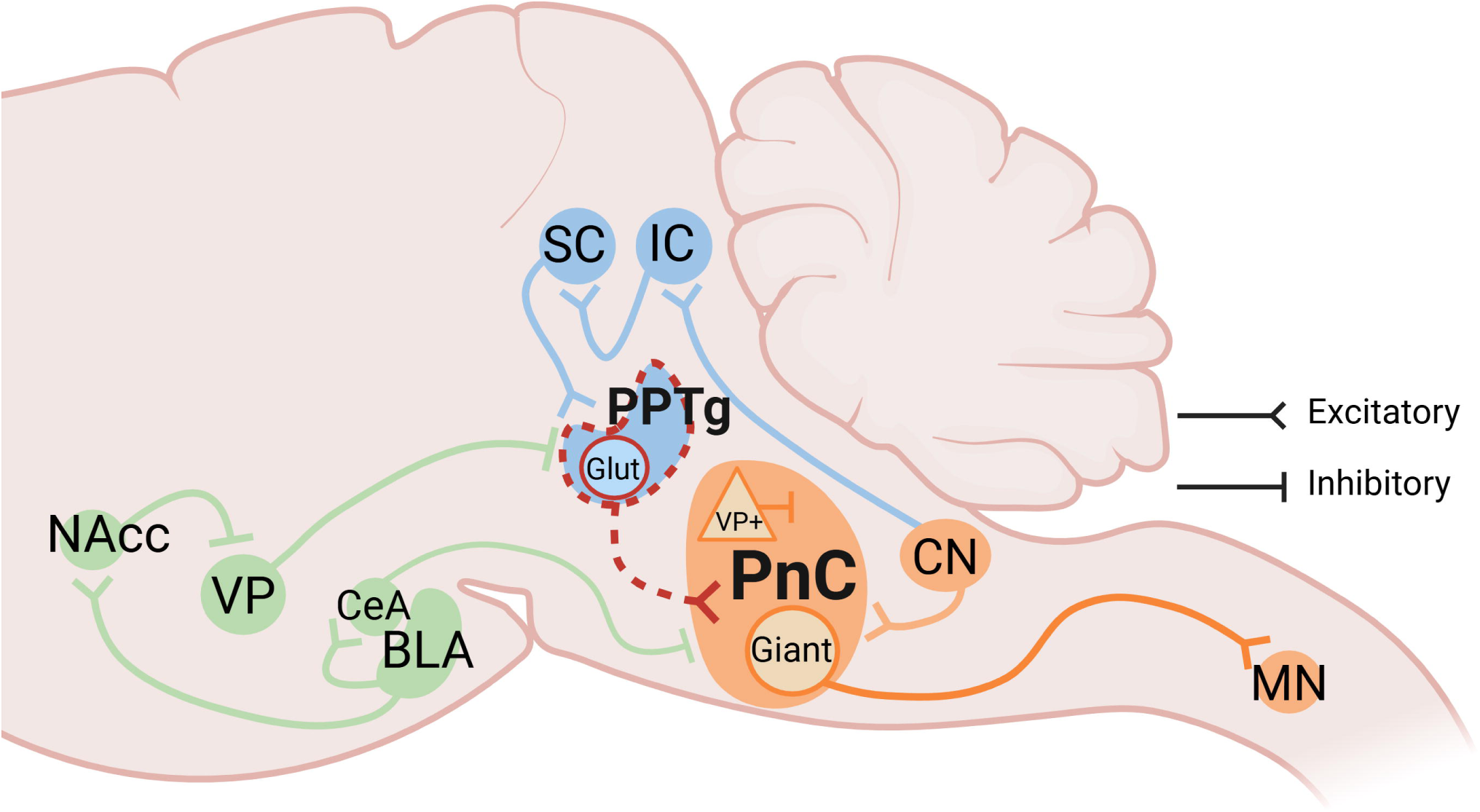
Proposed neural circuits involved in the acoustic startle response (ASR) and prepulse inhibition (PPI). The primary acoustic startle pathway (orange pathway) begins with auditory stimulation of the cochlear nuclei (CN), which then relays auditory signals to the giant neurons of the caudal pontine reticular nucleus (PnC) in the brainstem. PnC giant neurons directly excite spinal cord motor neurons (MN). The PPI pathway (blue pathway) responds to low-decibel acoustic prepulses and can inhibit a subsequent startle response via activation of the inferior colliculi (IC), superior colliculi (SC), and the pedunculopontine tegmental nucleus (PPTg). The PPI pathway is also modulated by midbrain and cortico-limbic structures (green pathway) comprised of the basolateral amygdala (BLA), the central amygdala (CeA), the nucleus accumbens (NAcc), and the ventral pallidum (VP). While these structures form the cortico-striato-pallido-pontine (CSPP) network, our study focuses on how PPTg glutamatergic neurons in particular modulate the PnC (dashed red) during startle and PPI.

The pedunculopontine tegmental nucleus (PPTg) is a midbrain region known to inhibit PnC-mediated startle responses. It was initially suggested that PPI requires PPTg cholinergic neuron activity. This was supported by studies showing that PPTg cytoarchitectural abnormalities are observed in patients with PPI deficits, that PPTg lesions reduce PPI, and that PPTg cholinergic inputs inhibit PnC giant neurons (Koch et al., 1993; Swerdlow and Geyer, 1993; Kodsi and Swerdlow, 1997; Bosch and Schmid, 2008). This initial hypothesis was recently ruled out by rat *in vivo* studies showing that optogenetic or chemogenetic manipulation of PPTg cholinergic neurons does not impact PPI (Azzopardi et al., 2018; Fulcher et al., 2020). Since the PPTg also comprises a substantial population of glutamatergic and GABAergic neurons (Wang and Morales, 2009; Luquin et al., 2018), these results suggest that non-cholinergic PPTg mechanisms play a crucial role in PPI. While the contribution of PPTg glutamatergic neurons to PPI remains unexplored, PnC-projecting PPTg glutamatergic neurons modulate locomotion in rodents (Martinez-Gonzalez et al., 2014; Huang et al., 2024b; Roseberry et al., 2016; Josset et al., 2018). Additionally, PPTg glutamatergic neurons can selectively target inhibitory striatal interneurons to enable feedforward inhibition (Assous et al., 2019).

We recently described a feedforward inhibitory mechanism by which glutamatergic neurons located in the central nucleus of the amygdala (CeA) activate GlyT2^+^ PnC glycinergic neurons in mice (Cano et al., 2021; Huang et al., 2024a). Our optogenetic experiments further confirmed that both CeA glutamatergic neurons and GlyT2^+^ PnC neurons contribute to PPI.

Based on these results showing that PnC-projecting CeA glutamatergic inputs can modulate startle and elicit PPI by activating PnC inhibitory neurons, here we tested the hypothesis that PPTg glutamatergic neurons also modulate PnC neuronal activity during startle and PPI. Using a combination of anatomical, *in vitro* optogenetic and electrophysiological approaches in mice, we report the existence of glutamatergic projections from the PPTg that innervate PnC giant and inhibitory neurons. We also functionally characterized PnC-projecting PPTg glutamatergic neurons using *in vivo* optogenetic manipulations during startle and PPI.

## Materials and Methods

All experimental procedures were performed in accordance with the National Institute of Health Guidelines, and were conducted in compliance with the UMass Institutional Animal Care and Use Committee (IACUC).

### Animals

Adult C57CBL/6J male and female mice aged between 5-6 months (Jackson Laboratories) were used for all procedures. Following surgery, animals were singly housed at the University of Massachusetts Amherst (UMass) vivarium with controlled temperatures and a 12-hour light/dark cycle with *ad libitum* access to food and water.

### Study design

To label the glutamatergic projections between the PPTg and PnC, we performed unilateral viral tracing experiments. For retrograde labeling experiments, mice (n=3) received unilateral injections of Fluorogold (FG; 2% in 9% Saline, Fluorochrome, Denver, CO) or retrograde virus AAVrg-hSyn-eGFP (Addgene #50465) in the **PnC**. For anterograde tracing, mice (n=8) were injected unilaterally with AAV-DJ-CaMKII-ChR2-mCherry (Addgene #26975) targeted at the **PPTg**. Two to five weeks later, we immunostained for the appropriate fluorescent reporter (eGFP or mCherry) along with either acetylchoinetransferase (ChAT) and GAD67 in the PPTg or postsynaptic density marker PSD-95, NeuN, and parvalbumin (PV) in the PnC.

To optogenetically inhibit PPTg glutamatergic axons in the PnC during behavior, we unilaterally injected WT mice (n=11) with AAV-DJ-CaMKIIα-NpHR3.0-eYFP (Addgene #26971) targeted at the **PPTg**. To optogenetically activate PPTg glutamatergic axons in the PnC during behavior, the same animals used for anterograde tracing were used for behavioral assessment. All mice undergoing behavioral assessment (n=19) received a fiberoptic implant ipsilateral to the injection site over the PnC. Behavioral experimental paradigms used a unilateral approach due to prior evidence that unilateral manipulations were sufficient to produce PPI deficits (Forcelli et al., 2012) and the clinical relevance demonstrating that unilaterally impacted brain regions can lead to PPI deficits (Sonnenberg et al., 2006). Four weeks following fiberoptic implant, animals were tested on acoustic startle response and prepulse inhibition behavioral assays in the absence and presence of yellow or blue light.

To investigate electrophysiological properties of PPTg glutamatergic neurons and their postsynaptic targets in the PnC, mice (n=10) received injections of AAV-DJ-CaMKII-ChR2-mCherry (Addgene #26975) targeted at the **PPTg**. After allowing for viral expression for 4-5 weeks, mice brains were collected for electrophysiology experiments.

### Surgery

Mice were sedated by inhaling 5% isoflurane vapors (Vedco, Saint Joseph, Missouri), placed in a stereotaxic apparatus, and immobilized using ear bars and a nose cone. Mice were maintained under anesthesia (1.5–3% isoflurane) for the duration of the surgery, and the head was leveled in all three axes. All injections were performed using a pressure micro-injector (Quintessential Stereotaxic injector, Stoelting Co., IL). Anterograde unilateral injections consisted of 0.3 µL total volume (2 injections of 0.15 µL each) targeted at the PPTg at the following coordinates relative to Bregma: **#1** AP: −4.84 mm, ML: +1.3 mm, DV: −3.75; and **#2** AP: −4.84 mm, ML: +1.3 mm, DV: −3.5mm. Retrograde unilateral injections consisted of 0.1 µL total volume aimed at the **PnC** (coordinates: AP: −5.4 mm, ML: +0.5 mm, DV: −5.15 mm; Paxino’s and Franklin, 2004). Following anterograde viral injection, a craniotomy was drilled ipsilateral to the injection site and dorsal to the implantation site, at the PnC level. A cannula guide with a 200-μm core optical fiber (Thorlabs, Newton, NJ) was then implanted over the PnC (coordinate: AP − 5.35mm, ML + 0.7mm, DV − 4.5 mm using StereoDrive Robot Stereotax by Neurostar), and fixed to the skull with dental cement (Parkell, Edgewood, NY). Following surgery, mice recovered for two-weeks, allowing for maximal transport of viral particles, or four-weeks to allow for opsin expression prior to behavioral testing.

### Behavior

Mice underwent the PPI task in a startle response system (PanLab System, Harvard Apparatus, Holliston, MA). Behavioral testing trials were designed, and data were recorded using PACKWIN V2.0 software (Harvard Apparatus, Holliston, MA). Sound pressure levels were calibrated using a standard SPL meter (model 407730, Extech, Nashua, NH). Mice were placed on a movement-sensitive platform. Vertical displacements of the platform induced by startle responses were converted into a voltage trace by a piezoelectric transducer located underneath the platform. Startle amplitude was measured as the peak maximum startle magnitude of the signal measured during a 1s window following the presentation of the acoustic stimulation. Prior to any testing session, animals were first handled and acclimatized to the testing chamber, where the mice were presented to a 65dB background noise, for 10 min. This acclimatization period was used to reduce the occurrence of movement and artifacts throughout testing trials. Following the acclimatization period, an input/output (I/O) assay was performed to test startle reactivity. This I/O test began with the presentation of a 40ms sound at different intensities (in dB: 70, 80, 90, 100, 110, and 120) every 15s, in a pseudorandomized order. Background noise (65dB) was presented during the 15s between sounds. A total of 35 trials (i.e., 7 sound intensities, each sound presented 5 times) were acquired and quantified. Startle reactivity, derived from this I/O assay, allowed the gain of the movement-sensitive platform to be set. This gain allowed the startle responses to be detected within a measurable range. Once determined, the gain for each experimental subject was kept constant throughout the remaining of the experiment. Startle habituation was effectively avoided by confirming that the startle response remained robust and intact in response to startle pulses presented prior to and throughout PPI protocols. PPI testing consisted of two different conditions as follows: (1) startling “pulse-alone” stimulations (120dB sound, 40ms, for baseline startle amplitude), and (2) combinations of a prepulse (75dB sound, 20ms) followed a 120dB startling pulse (40ms) at 8 different interstimulus intervals (ISIs; in ms): 10, 30, 50, 100, 200, 300, 500, and 1000 (end of prepulse to onset of startle pulse). The intertrial interval of these two conditions was 29s during which 65dB background noise was present.

For combined optogenetic manipulations and PPI testing, animals injected with viral vectors AAV-DJ-CamKIIa-NpHR3.0-eYFP (N = 11 mice) or AAV-DJ-CamKIIa-ChR2-mCherry (N = 8 mice) were tested in the startle chamber. These animals were closely monitored to ensure that they were comfortably tethered to an optic fiber, which exited through a small opening from the roof of the startle chamber. The optic fiber (200 μm diameter, Thorlabs, Newton, NJ) was connected to the animal’s head via a cannula implanted on the head of the mouse with a zirconia sleeve (Thorlabs, Newton, NJ). Animals were tethered ∼ 15min in the startle chamber to acclimate while 65 dB background noise was presented. Optogenetic stimulation was triggered by a signal from the Packwin software (PanLab System; Harvard Apparatus, Holliston, MA), which was transformed into a TTL pulse. This TTL pulse triggered a waveform generator (DG1022, Rigol Technologies), which was used to modulate light stimulation. Photo-stimulation used a blue 473-nm laser (Opto Engine LLC, Midvale, UT) for ChR2 activation, while photo-inhibition used a yellow 593.5-nm laser (Opto Engine LLC, Midvale, UT) for NpHR3.0 activation. During PPI trials paired with optogenetic inhibition or activation, a train of light stimulation (1ms light ON, 199ms light OFF) was delivered at 5 Hz and was either (1) delivered 400ms prior to and concurrent to the pulse-alone stimulation, or (2) delivered 400ms prior to and concurrent to the prepulse stimulation, lasting the entire interstimulus interval. During PPI trials with optical stimulation used as a prepulse, a 5Hz stimulation train (3 pulses of 15ms) was delivered at 400ms prior to and directly before various ISI (10, 30, 50, 100, 200, 300, 500, and 1000 ms) prior to the startling pulse. At the end of each experiment, histological analyses were performed to confirm that (1) the injected viral particles were confined to the PPTg, and (2) the cannula guide placement was successfully aimed at the PnC. If these criteria were not met, the subject was excluded from the study.

### In vitro electrophysiology

Brainstem slices for electrophysiological recordings were prepared using protocol described previously (Ting et al., 2018), with modifications. Mice (N = 20) were anesthetized in an induction chamber with 5% isoflurane in an oxygenated environment for 1 min or until completely unresponsive to toe pinch. Mice were removed from induction chamber, tested for lack of reflex response, and quickly decapitated. The head was submerged in chilled NMDG-HEPES aCSF (in mM): NMDG (92), KCl (2.5), NaH2PO4 (1.25), NaHCO3 (30), HEPES (20), glucose (25), thiourea (2), Na-ascorbate (5), Na-pyruvate (3), CaCl2 (0.5), and MgSO4 (10) (bubbled with 95 % O2 / 5 % CO2 and pH to 7.3–7.4 with HCl). The brain was exposed and swiftly removed from skull and submerged in chilled NMDG-HEPES aCSF, prior to sectioning of brainstem, mounting to platform using methyl-methacrylate glue, and submerging in chilled slicing chamber in Leica VT 1200s vibratome (Buffalo Grove, IL). Brainstem were sliced in coronal sections using zirconium ceramic injector blade (EF-INZ10, Cadence Blades, Staunton, VA). PnC and PPTg containing slices were collected and transferred to interface chamber containing slice recovery buffer, maintained at 35 °C, containing (mM): NaCl (92), KCl (2.5), NaH2PO4 (1.25), NaHCO3 (30), HEPES (20), glucose (25), thiourea (2), Na-ascorbate (5), Na-pyruvate (3), CaCl2 (0.5), and MgSO4 (10) (bubbled with 95 % O2 / 5 % CO2 and pH to 7.3–7.4 with NaOH). Slices were subject to “Na spike-in,” according to protocol and age, as previously described (Ting et al., 2018). Upon completion of Na-spike-in, slices were transferred to artificial cerebral spinal fluid (aCSF) maintained at room temperature (22 °C) containing (in mM): NaCl (124), KCl (2.5), NaH2PO4 (1), NaHCO3 (25), glucose (10), MgSO4 (1), CaCl2 (2) (bubbled in 95 % O2 / 5% CO2) for minimum of two hours prior to electrophysiological recordings.

Patch-clamp electrophysiological recordings were performed in brainstem slice preparations, under the whole-cell mode at room temperature with either EPC10/2 amplifier (HEKA Elektronic, Lambrecht, Germany) or Multiclamp 700B (Molecular Devices, San Jose, CA), and data were acquired with a PC computer running either Patchmaster (HEKA) or pClamp 10 (Molecular Devices), respective to amplifier. Brain slices were bathed in artificial cerebral spinal fluid (aCSF) maintained at room temperature (22 °C) containing (in mM): NaCl (124), KCl (2.5), NaH2PO4 (1), NaHCO3 (25), glucose (10), MgSO4 (1), CaCl2 (2), bubbled in 95 % O2 / 5% CO2. Recording chamber containing brain slice were gravity fed ACSF at a rate of 2-3 ml/minute. Cells were impaled with patch pipettes pulled from borosilicate glass capillaries (1B150F-4, World Precision Instruments, Sarasota, FL) using P-97 horizontal pipette puller (Sutter Instruments, Novato, California). The resistance of patch pipettes was typically 3-6 MΩ when filled with an internal solution containing (in mM): K-Gluconate (30), KCl, 10 HEPES (4), EGTA (0.1), phosphocrea-tine-Na2 (10), MgATP (4), Na2-GTP (0.3), and biocytin (10). The pH was adjusted to 7.35 with 1M KOH and the osmolality to 285–290mOsm. Neurons were compensated for capacitance and series resistance online to 90% using software driven capacitance control circuit. In voltage clamp experiments cells were held at −70mV to record spontaneous activity and during optogenetic stimulation. For current clamp experiments, cells were rejected if current compensation to maintain −70mV holding voltage exceeded 200pA.

In the PPTg, neurons expressing channelrhodopsin were targeted for patch clamp recordings, which were visualized by coexpression of the eYFP fluorescent marker. In the PnC, mCherry-fluorescent axon fibers from PPTg Glutamatergic neurons transfected with AAV-CaMKII-ChR2-mCherry were targeted by photo-stimulations for post-synaptic patch clamp recordings. PPTg or PnC neurons that were patched were allowed to equilibrate with pipette solution for minimum of 15 minutes prior to optogenetic stimulation. Optogenetic stimulation of channelrhodopsin-expressing PPTg axons were stimulated by TTL pulse to Plexon, PlexBright LD-1 single channel LED driver (Plexon, Dallas, TX) to activate an LED module (470 nm; Thor Labs) channeled to the tissue by fiber optic cable, centered over the patched cell using a micromanipulator. For single pulse and paired pulse stimulation, light pulse duration did not exceed 1ms. Light pulse train stimulation templates were generated in Igor Pro 6.0 (WaveMetrics, Lake Oswego, OR) through template feature in Patchmaster. Data were analyzed with either Igor Pro 6.0 or Clampfit (Molecular Devices).

### Immunohistochemistry

24-hours following the conclusion of behavioral testing, mice (N = 19) were anesthetized with isoflurane and exsanguinated by transcardial perfusion with approximately 50mL chilled 0.9% saline solution followed by chilled 4.0% paraformaldehyde (PFA) solution in phosphate buffered saline (PBS). After perfusion, animals were decapitated and the brains were extracted and harvested followed by post-fixing in PFA overnight at 4°C for approximately 12-18 hours and cryoprotection in 30% sucrose at 4°C for approximately 36-48 hours. The brains were drained of excess sucrose, and frozen in super cooled hexanes and stored at −80°C. Then, the brains were sectioned on a Leica CM3050 S Cryostat (Leica, Wetzlar, Germany). The sections were cut into 1-in-6 series throughout the extent of the PnC and PPTg at a thickness of 30µm in the coronal plane of section to survey the whole area in which the PPTg and PnC areas are present, since the areas are in a short distance from one another. Sections were stored in cryoprotectant solution (50% 0.05M phosphate buffer, 3-& ethylene glycol, 20% glycerol) at ^-^20°C - ^-^80°C until stained and/or plated for visualization.

The viral constructs contained the reporter gene mCherry or eYFP which are translated into fluorescent proteins which were enhanced via immunohistochemistry for optimal visualization/labelling (chicken anti-mCherry, 1:1000, Abcam, ab205402; or chicken anti-GFP, 1:1000, Abcam, ab13970). Additional antibodies were used to double-label cells expressing specific proteins to reveal their identities. Fluorogold was not included since it already expresses fluorescent properties and does not require antibodies for visualization. The brain slices were collected and separated into two groups depending on whether they contained the PPTg or the PnC. The brain sections were rinsed off of cryoprotectant solution for 3 times for 5 minutes each using Tris-buffered saline (TBS; pH 7.4 at room temperature) and incubated in Blocking solution (NDS) (3% normal donkey serum; EMD-Millipore, Billerica, MA; catalog #S30-100ML; lot NG1827420 and 0.1% Triton X-100; Sigma-Aldrich, St. Louis, MO; catalog #T8532 in TBS) for 1 hour at room temperature. The PPTg sections were treated with goat anti-ChAT antibody (1:200; Millipore, AB144P) since labeling PPTg Cholinergic neurons still constitutes the best way to delineate its borders. The PnC sections were treated with either a guinea pig anti-NeuN antibody (1:1000; Sigma Aldrich, ABN90P), a marker of mature neurons, or a rabbit anti-parvalbumin antibody (1:1000; Abcam, ab11427), a marker of inhibitory neurons, and a goat anti-PSD95 antibody (1:500; Abcam, ab12093), a marker of excitatory synapses. Slices were incubated in primary antibodies for 1-2 days at 4°C. The slices were then washed of the primary antibody in TBS for 3 times for 10 minutes each at room temperature and incubated in combinations of secondary antibodies used at a concentration of 1:500 (donkey anti-guinea pig DyLight405, Jackson IRL, 706-475-148; donkey anti-goat Cy3 or AF488, Jackson IRL; donkey anti-rabbit DyLight405, Jackson IRL, 711-475-152; donkey anti-chicken Cy3 or AF488, Jackson IRL) for 3-4 hours at room temperature, then rinsed again. The trays with the tissue were covered by aluminum foil to prevent photo-bleaching during the antibody treatment. The slices were then mounted on super frost slices, air-dried and cover-slipped.

### Statistical analysis

Cell counting of fluorescent or Fluorogold^+^ somata within the PPTg or PnC was performed in a tissue slice series of 6 slices. Statistical analyses were performed using SigmaPlot (Systat Software, Inc., San Jose, CA) and GraphPad Prism (GraphPad Software, La Jolla, CA). Normality and equal variance of the data were first tested, and data transformations were made before performing further statistical analyses. We determined the significance of the interaction between the factors assessed using ANOVA. For the results of whole-cell patch clamp recordings with receptor antagonists, one-way repeated measures (RM) ANOVA and Tukey post hoc testing were used to assess the effect of the receptor antagonists on the light-evoked events. PPI was defined and measured as [1–(startle amplitude during “Prepulse+Pulse” trials/startle amplitude during “Pulse” trials)] × 100. Two-way Repeated Measures (RM) ANOVA was used to assess the effect of the vector used, light, sound intensity/inter-stimulus interval (ISI), and light interaction and the interaction among groups. For optical stimulation experiments where the photo stimulation of neurons fibers was used as a prepulse *in vivo*, two-way RM ANOVA was used to assess the effect of the stimulation, ISI, ISI and stimulation interaction and interaction among groups. Then, Tukey testing was applied for post hoc comparisons. A confidence level of p < 0.05 was considered statistically significant. Sample sizes were chosen based on expected outcomes, variances, and power analyses. Data are presented as means ± SEM. N indicates total number of animals; n indicates total number of brain slices or testing trials. Adobe Illustrator and BioRender were used to create figures.

## Results

### PPTg glutamatergic neurons innervate the PnC

To quantify the number of putative PPTg glutamatergic neurons that project to the PnC, we first injected a retrograde AAV that transduced the fluorescent reporter TdTomato under the control of the CAG promoter (AAVrg-CAG-tdTomato) into the PnC of WT mice (N = 3). As previously reported in rats (Wang and Morales, 2009; Luquin et al., 2018), we identified retrogradely labeled tdTomato^+^ PPTg neurons that were ChAT-immunonegative in mice (ChAT^-^; **Suppl. Fig. 1**). Such ChAT^-^ PnC-projecting PPTg neurons are likely glutamatergic since they were labeled with tdTomato but not co-labeled with the inhibitory neuron GAD67 antibody (**Suppl. Fig. 1D**).

Using *in situ* hybridization, we previously showed that CamKIIα is exclusively expressed by PPTg glutamatergic neurons (Cano et al., 2021). Therefore, to track how PPTg glutamatergic fibers course within the PnC, we took an anterograde approach and successfully targeted CamKIIα^+^PPTg neurons by injecting AAV-DJ-CamKIIα-eYFP (**Fig. 2**) in the PPTg of WT mice (N=8). Following PPTg injection (**Fig. 2 B; also see Suppl. Fig. 2**), mCherry^+^ PPTg glutamatergic fibers were seen across both ventrolateral and ventromedial areas of the PnC delineated by the 7th cranial nerve (**Fig. 2 C, D**). Interestingly, PPTg glutamatergic innervation in the PnC appears denser on the side ipsilateral to the PPTg viral injection (**Suppl. Fig. 1 C, D**). We then used immunohistochemistry in combination with confocal microscopy to evaluate the apposition between mCherry^+^ PPTg glutamatergic fibers and PnC sections stained for PSD95, an adaptor protein involved in clustering postsynaptic receptors at glutamatergic synapses (Cho et al., 1992; Kornau et al., 1995) in the entire extent of the PnC (**Fig. 2 D**). Our volume rendering results and angular sectioning of PSD-95^+^ putative synaptic contacts suggest that the PPTg sends glutamatergic projections that form glutamatergic synapses with postsynaptic PnC neurons located ventrolaterally and ventromedially.

**Figure 2.**
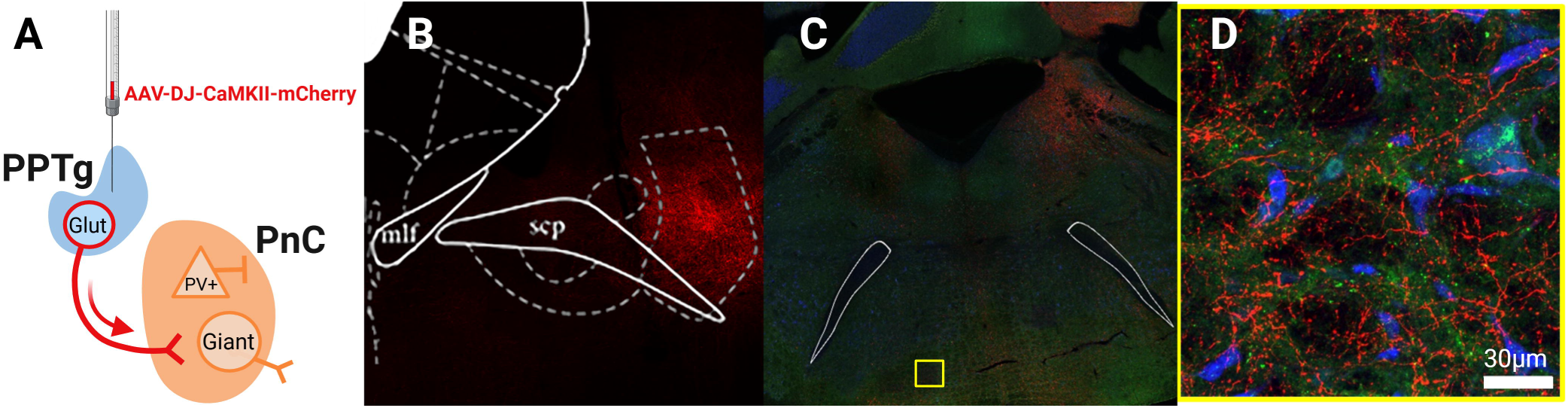
Anterograde labeling of PnC-projecting PPTg glutamatergic neurons. (**A**) Schematic illustrating the unilateral injection of AAV-DJ-CaMKIIα-mCherry in the PPTg of wildtype mice. (**B**) Representative AAV microinjection in the PPTg at 20x magnification. (**C**) Labeled PPTg glutamatergic axons fibers (red) in the PnC at 20x magnification. (**D**) Close up of the region delineated with the yellow box in (C), showing glutamatergic axons (red) closely apposed to PnC neurons labeled with the nucleic marker NeuN (blue) and the excitatory post-synaptic marker PSD95 (green) at 40x magnification. Images are representative of N = 8 mice. Scale bar is 30 μm in (D).

### Silencing PPTg-PnC glutamatergic synapses enhances PPI without affecting baseline startle

Next, we aimed to determine the contribution of PPTg glutamatergic neurons projecting to the PnC during acoustic startle responses and PPI, *in vivo*. To do so, we conducted optogenetic experiments to silence PPTg-PnC glutamatergic synapses in WT mice by transducing PPTg glutamatergic neurons with the yellow-light sensitive inhibitory optogenetic tool Halorhodopsin (NpHR3.0). Mice were injected with the AAV-DJ-CamKIIα-NpHR3.0-eYFP viral vector unilaterally in the PPTg. Following the intracranial viral injection, an optic fiber was implanted ipsilateral to the injection site at the level of the PnC, to photo-inhibit PPTg fibers/terminals expressing NpHR3.0 with yellow light (**Fig. 3 A;** N = 11).

**Figure 3.**
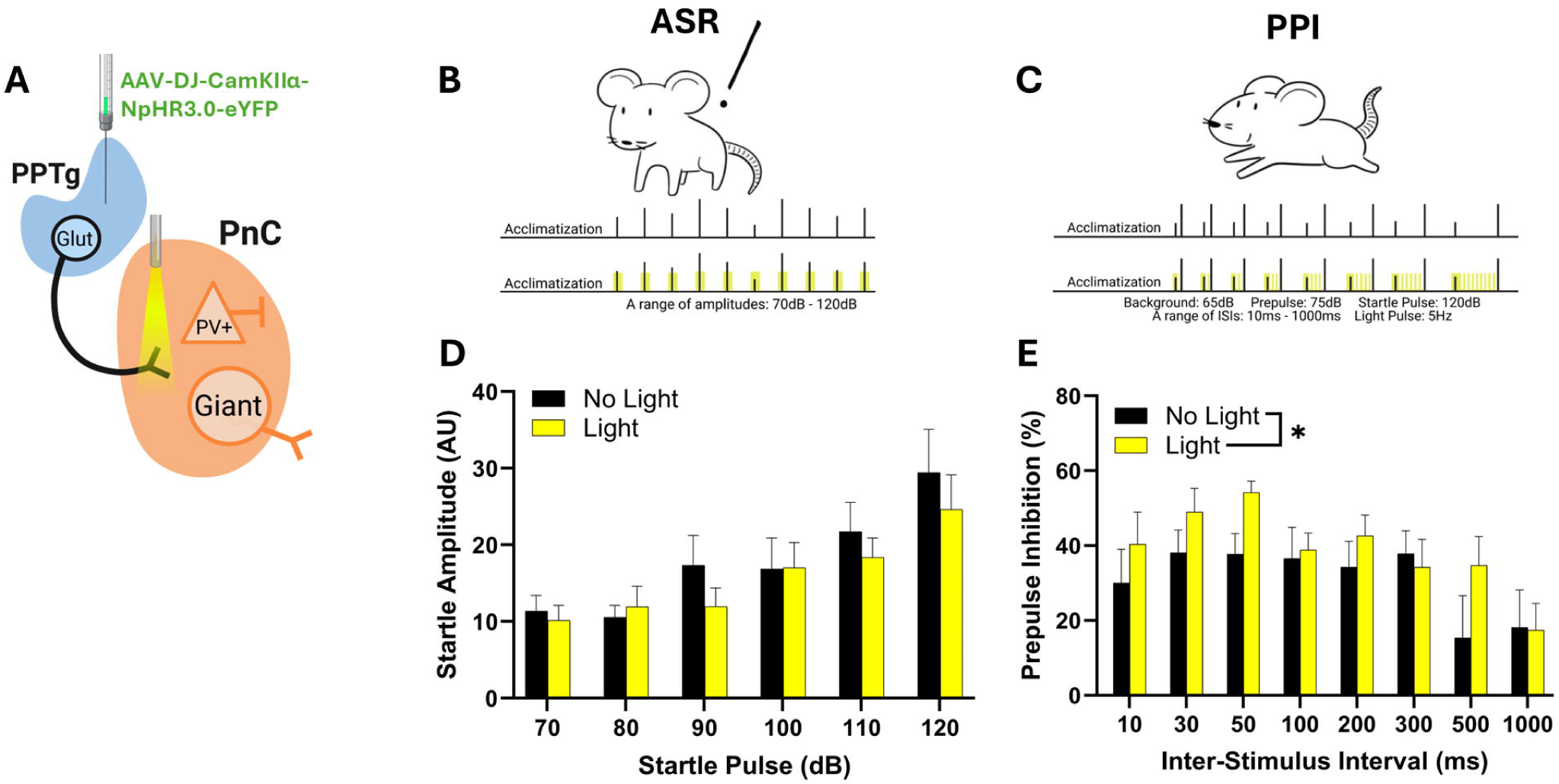
Silencing PPTg-PnC glutamatergic synapses increases PPI without affecting baseline startle. (**A**) Schematic illustrating the location of AAV injection of Halorhodopsin (NpHR3.0) and yellow light delivery for the optogenetic silencing of PPTg-PnC glutamatergic synapses. (**B**) Schematic illustrating the acoustic startle reflex protocol performed using mice injected with Halorhodopsin, in the absence (top) or presence (bottom) of yellow light. (**C**) Schematic illustrating the PPI protocol performed using mice injected with Halorhodopsin, in the absence (top) or presence (bottom) of yellow light. (**D**) Graph showing that silencing PPTg-PnC glutamatergic synapses with yellow light did not affect basal startle amplitude elicited by 70–120 dB acoustic pulses (Two-Way RM ANOVA: F(1) = 0.7178, p = 0.4167) (**E**) Graph showing that silencing PPTg-PnC glutamatergic synapses with yellow light significantly increased PPI when delivered during acoustic prepulses and inter-stimulus intervals (ISIs) (Two-Way RM ANOVA: F (1, 87) = 5.968, p = 0.0166). N = 11 mice. Data are represented as mean ± SEM. *p < 0.05

To determine if PPTg-PnC glutamatergic synapses contribute to baseline acoustic startle responses, we assessed how photo-inhibiting PPTg-PnC glutamatergic synapses with yellow light alters startle reactivity at increasing sound levels in mice expressing NpHR3.0 in PPTg glutamatergic neurons. In these mice, in the absence of yellow light photo-inhibition, sound intensities of 90 dB and beyond elicited an acoustic startle reflex (**Fig. 3 B, D**), measured as an involuntary innate defensive response occurring within milliseconds after the onset of startling stimulus and characterized by a whole-body flexor muscle contraction (Davis et al., 1982; Lee et al., 1996; Swerdlow et al., 1999). Interestingly, photo-inhibition of PPTg-PnC glutamatergic synapses concurrent with such acoustic startling stimuli did not affect startle amplitude (**Fig. 3 D**; [Two-Way RM ANOVA, light: (F(1, 10) = 0.7178, p = 0.4167); sound intensity × light interaction: (F(5,50) = 0.9854, p = 0.436)). These data confirm that PPTg glutamatergic neurons projecting to the PnC do not contribute to baseline startle reactivity.

We then tested whether silencing the PPTg-PnC glutamatergic synapses alters PPI *in vivo*. To do so, we photo-inhibited this pathway starting from the beginning of the prepulse until the end of the interstimulus intervals (**Fig. 3 C, E**). Photo-inhibition of PPTg-PnC glutamatergic synapses significantly increased PPI (**Fig. 3 E**; Two-Way RM ANOVA: p=0.0166). These photo-inhibition results suggest that PPTg glutamatergic neurons provide an excitatory drive to the PnC startle circuit during PPI.

### Activating PPTg-PnC glutamatergic synapses modulates acoustic startle and PPI

Next, we tested whether photo-activating PPTg-PnC glutamatergic synapses concurrent with an acoustic prepulse impacts baseline startle (**Fig. 4**). To do so, WT mice were injected with the blue-light sensitive excitatory optogenetic vector AAV-DJ-CamKIIα-ChR2-mCherry in the PPTg and implanted with an optic fiber positioned above the PnC. In these mice, PPTg-PnC glutamatergic synapses were photo-activated with short trains of blue light at 5Hz, concurrent with an acoustic startling (pulse-alone) stimulation. Photo-activation of ChR2^+^ PPTg-PnC glutamatergic synapses concurrent with acoustic startling stimulations did not significantly impact baseline startle (**Fig. 4 E**; Two-Way RM ANOVA: p = 0.1192).

**Figure 4.**
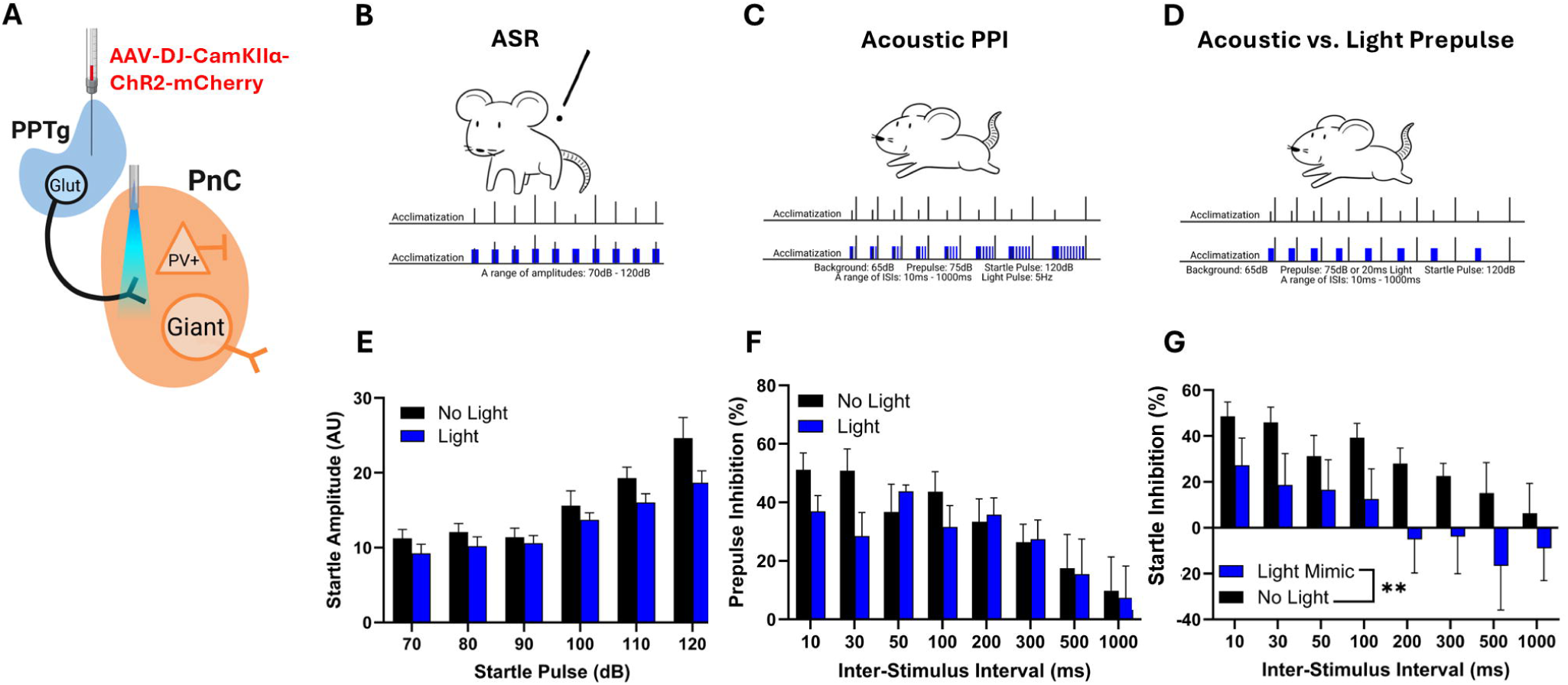
Photo-stimulation of PPTg-PnC excitatory synapses alone can modulate startle. (**A**) Schematic illustrating the location of AAV injection of Channelrhodopsin-2 (ChR2) and blue light delivery for the optogenetic activation of PPTg-PnC glutamatergic synapses. (**B**) Schematic illustrating the acoustic startle reflex protocol performed using mice injected with ChR2, in the absence (top) or presence (bottom) of blue light prior to and during 70–120 dB acoustic pulses. (**C**) Schematic of PPI protocol performed using mice injected with ChR2, in the absence (top) or presence (bottom) of blue light paired with acoustic prepulses and inter-stimulus intervals (ISIs). (**D**) Schematic illustrating the PPI Light mimic protocol performed using mice injected with ChR2, where blue light was present in lieu of an acoustic prepulse at various ISIs. (**E**) Graph showing no significant impact on startle response in the presence of blue light (Two-Way RM ANOVA: F (1, 6) = 3.299, p = 0.1192). (**F**) Graph showing no significant differences in PPI in the presence of blue light (Two-Way RM ANOVA: F (1, 6) = 0.3677, p=0.5665). (**G**) Graph showing that blue light significantly decreased startle at ISIs ≤100ms and potentiated startle response at ISIs > 100ms (Two-Way RM ANOVA: F (1, 47) = 26.40, p<0.0001). N = 7 mice per group. Data are represented as mean ± SEM. *p < 0.05, **p < 0.01

Then, to test whether stimulating this glutamatergic connection could alter PPI, PPTg-PnC glutamatergic synapses were photo-activated during PPI using blue-light. Photo-activation started at the beginning of the prepulse and lasted until the end of the interstimulus intervals (**Fig. 4 C, F**). Our results show that photo-activating PPTg-PnC glutamatergic synapses did not alter PPI (**Fig. 4 F**; Two-Way RM ANOVA, *p*=0.56). These results suggest that artificially activating PPTg glutamatergic neurons does not further alter acoustic PPI.

We next aimed to test whether PPTg-PnC glutamatergic synapses alone are sufficient to modulate subsequent baseline startle responses (**Fig. 4 D, G**). To do so, we assessed the effect of photo-activating PPTg-PnC glutamatergic synapses alone, prior to an acoustic startling stimulation, in the absence of an acoustic prepulse. PPTg-PnC glutamatergic synapses were photo-activated with a 20ms-blue light pulse prior to the startling stimulation, at intervals relevant to acoustic PPI (**Fig. 4 D, G**). Interestingly, our results show that photo-activating PPTg-PnC glutamatergic synapses in the absence of an acoustic prepulse had two effects: **1)** it reduced subsequent startle responses, mimicking an acoustic prepulse <u>inhibition</u> effect (or PPI) at *short* ISIs i.e., ≤100ms, but **2)** it enhanced subsequent startle responses, mimicking an acoustic prepulse <u>facilitation</u> effect (potentiation or “negative PPI”) at *longer* ISIs i.e., >100ms (**Fig. 4 G**; Two-Way RM ANOVA: p <0.0001). These results suggest that PPTg glutamatergic neurons are sufficient to bi-directionally modulate subsequent startle responses possibly by providing an excitatory drive to the startle circuit and activating inhibitory or excitatory elements within the PnC, depending on the timing of their activation.

### Electrophysiological properties of PPTg glutamatergic neurons

Before further investigating how PPTg glutamatergic neurons modulate the PnC startle circuit, we took advantage of the fact that CamKIIα-mCherry^+^ PPTg neurons expressing ChR2 are sensitive to blue light photo-stimulation to document their basic electrical properties (N = 10 mice; n = 10 slices and 26 neurons). To do so, we performed *in vitro* patch clamp recordings in acute PPTg slices of WT mice previously injected with AAVDJ-CamKIIα-ChR2-mCherry in the PPTg. This transduced PPTg glutamatergic cells with ChR2 and mCherry (**Suppl. Fig. 3**). From these ChR2-mCherry^+^ PPTg glutamatergic neurons held at −70 mV, spontaneous excitatory post-synaptic currents (sEPSC) with a mean amplitude of 14.1 ± 2 pA were recorded and not significantly different than the sEPSC amplitude of neighboring ChR2-mCherry^-^ PPTg neurons (mCherry^-^ −18.46 ± 1.85 pA, mCherry^+^ −16.55 ± 1.31 pA; T-test, *p* = 0.4059; **Suppl. Fig. 3 A, B**). While the half width of the action potentials did not differ between cell types (**Suppl. Fig. 3 B, *Inset;*** mCherry^+^: 1.41 ± 0.164 ms vs. mCherry^-^: 1.62 ± 0.265 ms; T-test, *p* = 0.4812), current injections elicited action potentials firing at a maximum rate of 9.4 ± 1.91 Hz in ChR2-mCherry^+^ PPTg glutamatergic neurons vs. 12.4 ± 1.01 Hz in ChR2-mCherry^-^ PPTg neurons (**Suppl. Fig. 3 C**). Other intrinsic properties such as input resistance, I/V curves and rheobase were not different.

Importantly, photo-stimulation induced large current responses in ChR2-mCherry^+^ PPTg neurons (**Fig. 5 A**), confirming that our stimulation protocol can successfully activate these PPTg glutamatergic neurons. Next, to quantify the blue-light sensitivity of ChR2-mCherry^+^ PPTg glutamatergic neurons, we recorded their activation pattern in response to short photo-stimulation trains consisting of blue light pulses of 0.1ms, 0.3ms and 1.0ms in duration, applied at 10, 20 and 40 Hz (**Fig. 5 B, C**). Photo-stimulation trains using 0.1ms light pulses were sufficient to elicit few action potentials but inefficient at eliciting a current response in ChR2-mCherry^+^ PPTg neurons, at the three frequencies tested. In contrast, photo-stimulation trains using 0.3ms and 1.0ms pulses induced sustained firing and current responses at all three frequencies (**Fig. 5 B, C**). These optically-induced current responses reliably followed the stimulation frequency, and the cumulative charge was correlated to the pulse duration (**Fig. 5 D**).

**Figure 5.**
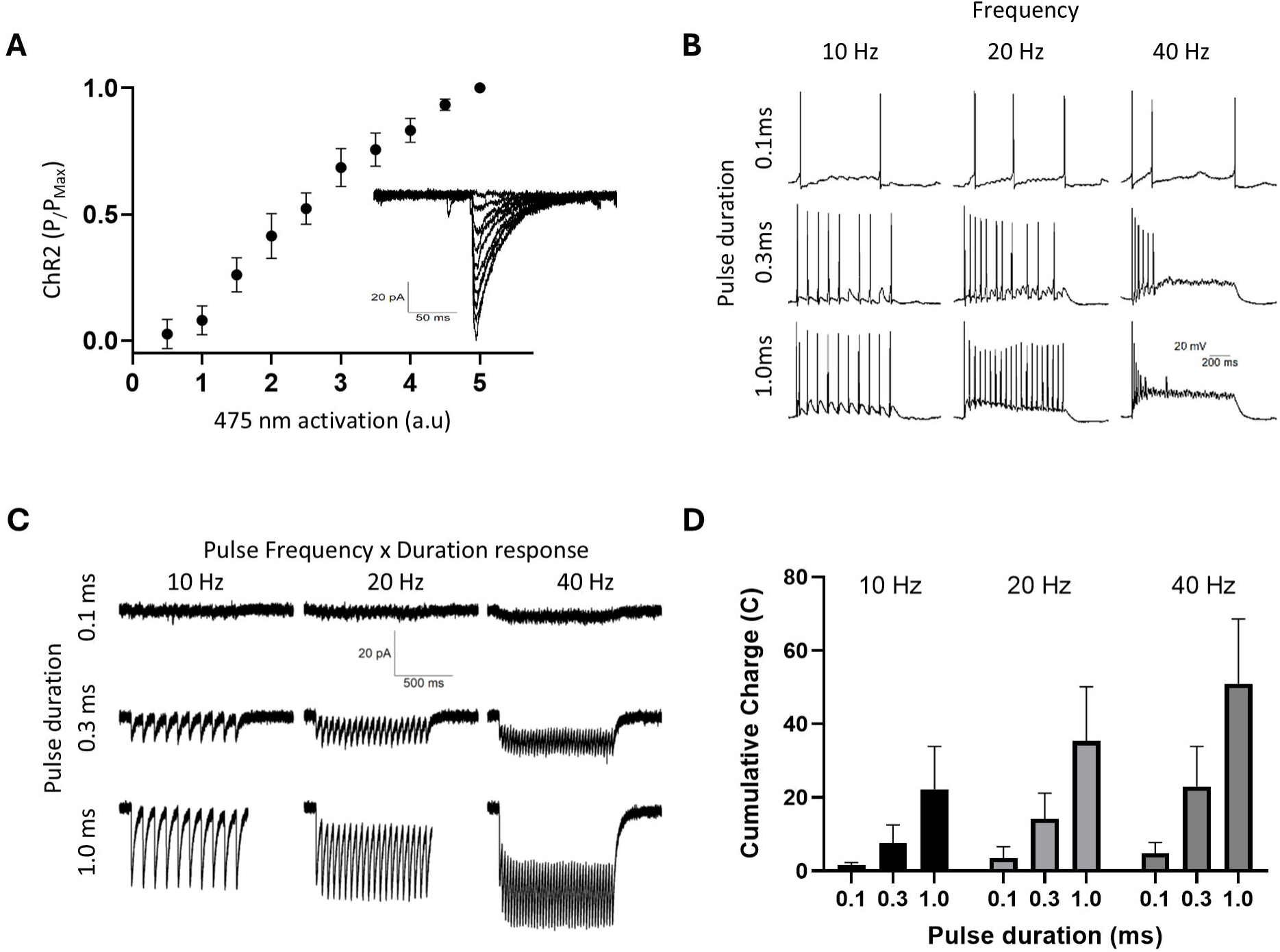
Light-evoked responses recorded in CamKIIα-ChR2-mCherry^+^ PPTg neurons. **(A)** Graph showing the mean EPSC expressed as fraction of maximal peak, at increasing blue-light photo-stimulation intensity (arbitrary units). Inset Representative excitatory post-synaptic current (EPSC) traces from a whole cell patched CamKIIα-ChR2-mCherry^+^ PPTg neuron responding to 1 ms blue light photo-stimulation of increasing intensity. **(B)** Sample traces showing blue light-evoked firing pattern recorded in CamKIIα-ChR2-mCherry^+^ PPTg neurons, in response to trains of light photo-stimulation of 0.1, 0.3 and 1.0 ms in duration and at 10 Hz, 20 Hz and 40 Hz stimulation frequencies. **(C)** Sample traces showing blue light-evoked EPSCs recorded in CamKIIα-ChR2-mCherry PPTg^+^ neurons, in response to trains of light photo-stimulation of 0.1, 0.3 and 1.0 ms in duration and at 10 Hz, 20 Hz and 40 Hz stimulation frequencies. **(D)** Plot showing the mean cumulative charge (integral of current over time) of EPSCs during the stimulation trains shown in (C). Within each frequency 0.1 ms blue light pulses elicited the smallest EPSCs, followed by 0.3 ms pulses, while 1.0 ms blue light pulses elicited the largest EPSCs for each group. N = 10 mice, n = 10 slices. Data are represented as mean ± SEM.

Next, we wanted to determine whether the optically evoked currents recorded in ChR2-mCherry^+^ PPTg neurons were synaptic in nature or resulted from activating non-synaptic membrane channels located on the soma of these ChR2-mCherry^+^ PPTg neurons. To test the possibility that these photocurrents are synaptic, we delivered pairs of light pulses, using standard short-term synaptic plasticity stimulation protocols (Zucker and Regehr, 2002). We hypothesized that if these currents are synaptic, then synapses with low neurotransmitter release probability (*p*) would exhibit short-term facilitation and a paired-pulse current ratio>1, whereas synapses with high *p* would display synaptic depression and a paired-pulse current ratio<1. Interestingly, our results show that within PPTg slices, optically stimulating ChR2-mCherry^+^ PPTg glutamatergic cells with pairs of light pulses elicited a paired-pulse ratio>1 at interstimulus intervals (ISIs) shorter than 50ms, for both voltage clamp (EPSCs) or current clamp (i.e., EPSPs) (**Suppl. Fig. 3 D-F**). At all the other intervals tested, the paired-pulse ratio was equal to 1, suggesting no plasticity.

To further test the synaptic nature of the light-induced paired responses recorded in ChR2-mCherry^+^ PPTg neurons, paired photo-stimulations were delivered after the bath-application of the glutamate receptor blockers, CNQX and APV. The pharmacological blockade of glutamate receptors did not affect the responses recorded in PPTg neurons or the paired-pulse ratios, indicating that the electrical activity of ChR2-mCherry^+^ PPTg neurons results from light-activated membrane channels rather than from synaptic activity between neighboring PPTg neurons (**Suppl. Fig. 3 D-F;** RM ANOVA, *p*>0.05). These findings further suggest that PPTg glutamatergic neurons function as projection neurons, sending inputs outside the PPTg without forming local synaptic connections within the PPTg.

### PPTg glutamatergic neurons activate PnC giant neurons

Next, we aimed to investigate how PPTg glutamatergic neurons activate PnC neurons within the startle circuit. To do so, we injected the AAV viral vector AAV-DJ-CamKIIα-mCherry into the PPTg of WT mice. Post-hoc immunohistochemical analysis confirmed that the PPTg was successfully targeted. ChR2-mCherry^+^ PPTg glutamatergic fibers were observed in acute PnC slices obtained from the same mice (**Fig. 6 D**). Using these PnC slices, we then wanted to determine whether photo-stimulation of ChR2-mCherry^+^ PPTg glutamatergic fibers could elicit excitatory synaptic responses in PnC giant neurons identified by their size >35um (**Fig. 6 C**). These neurons displayed spontaneous currents of −23.33± 3.6 pA in amplitude and spontaneous action potentials with ½ width of 0.91 ± 0.185 ms. In these giant PnC neurons held at −70 mV, blue light photo-stimulation pulses evoked large currents (**Fig. 7 A**), as well as excitatory post-synaptic potentials (**Fig. 7 B**, *top traces*). These excitatory responses showed facilitation at short ISI (**Fig. 7 B**, *middle and bottom traces*) (EPSP PPR: 50ms = 1.6, 100ms = 1.2; EPSC PPR: 50ms = 1.70, 100 ms = 1.17). These results confirm that PPTg glutamatergic neurons establish excitatory synapses with PnC giant neurons and that they can display facilitation.

**Figure 6.**
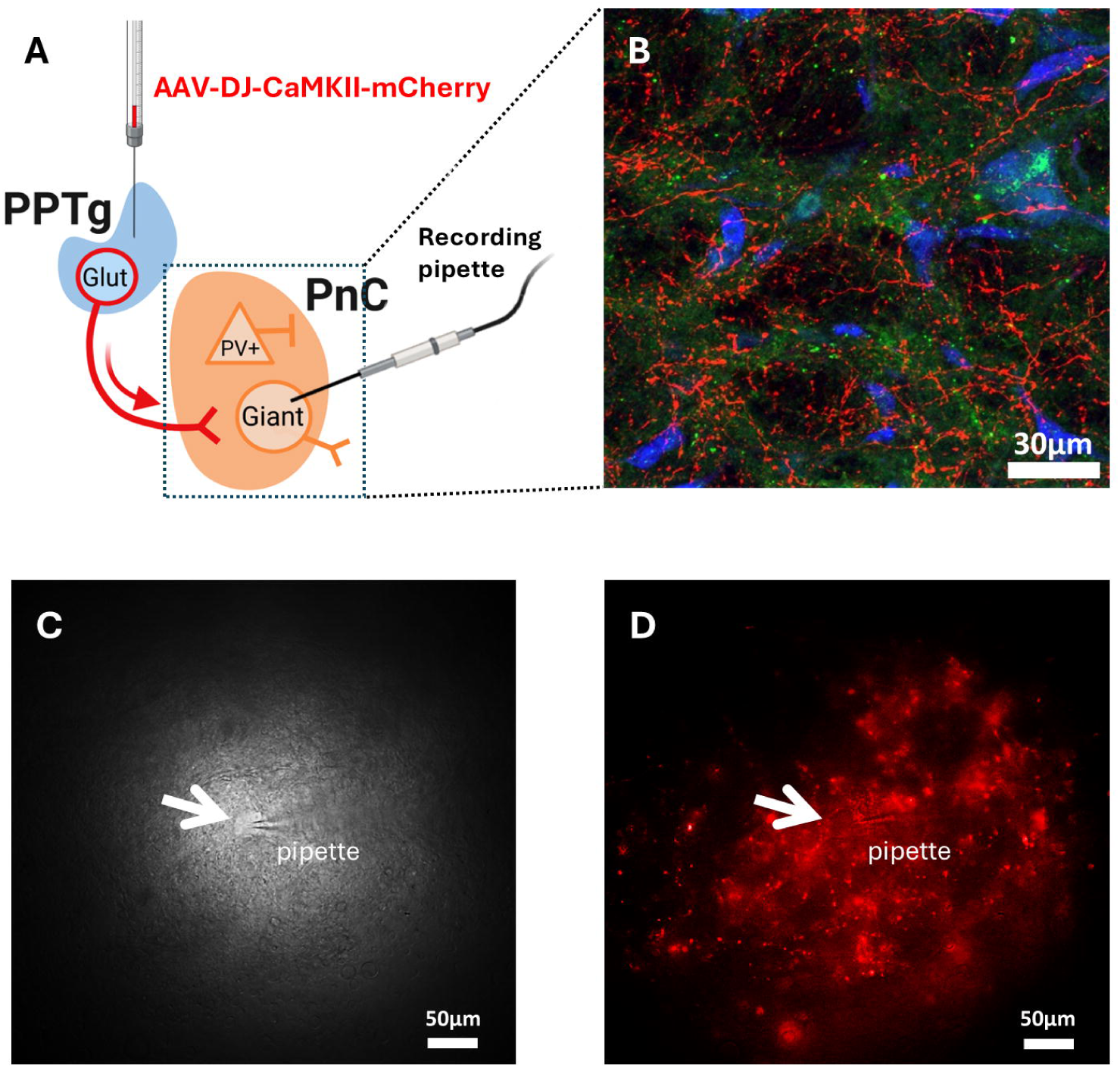
PPTg glutamatergic fibers innervate PnC giant neurons. **(A)** Scheme of the in vitro patch clamp recording experiment showing the blue-light photo-stimulation CamKIIα-ChR2-mCherry^+^ PPTg fibers and whole cell recording of giant neurons in PnC sections of WT mice. **(B)** Representative image of CamKIIα-ChR2-mCherry^+^ PPTg fibers (red) coursing within a PnC coronal section stained with the excitatory postsynaptic marker PSD95 (green) and the nuclei marker DAPI (blue). **(C, D)** Image of a patched PnC neuron (white arrow) innervated by CamKIIα-ChR2-mCherry^+^ PPTg fibers. Representative of N = 10 mice. Scale bars: B = 30μm; C, D = 50μm

**Figure 7.**
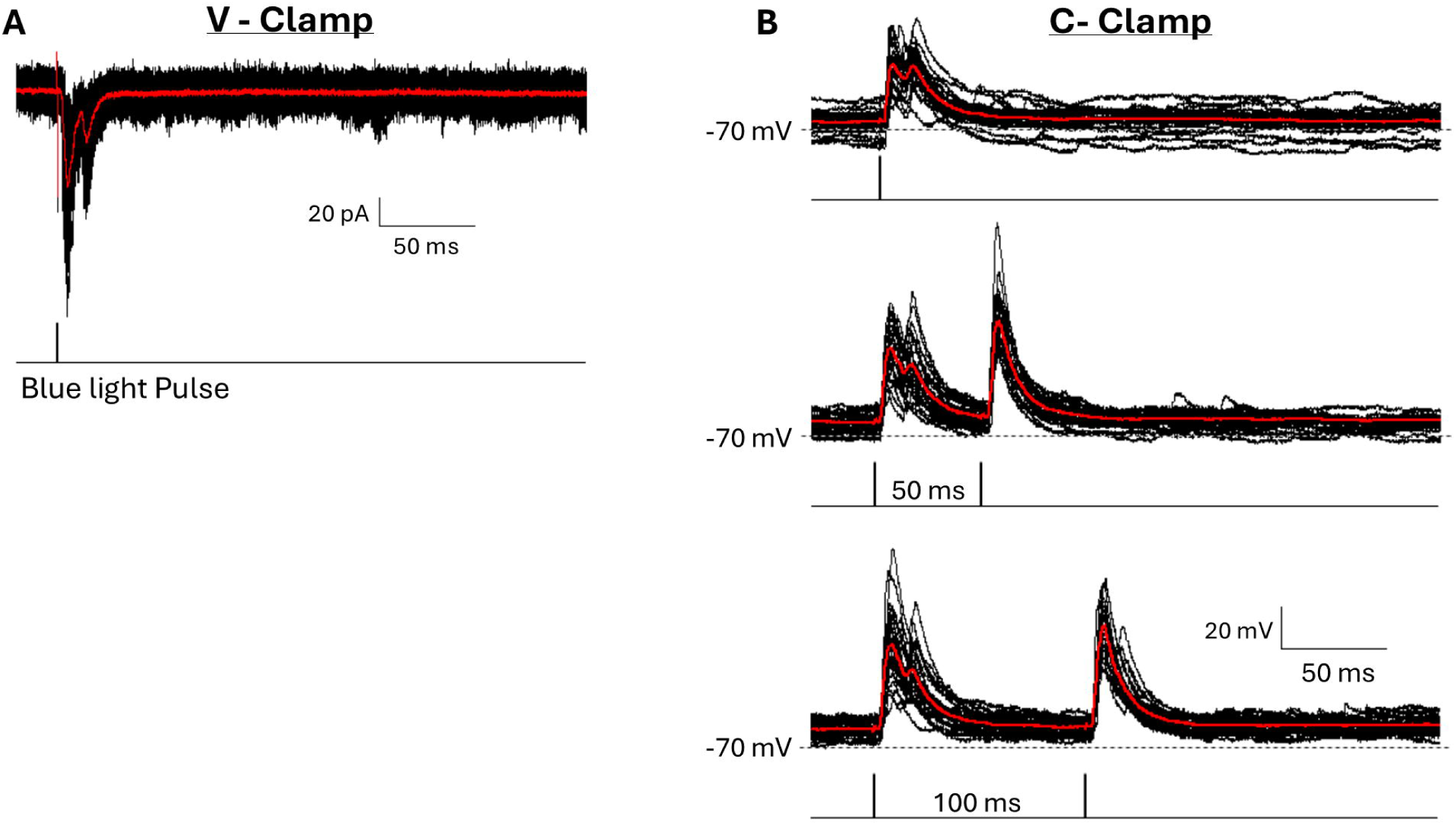
Photo-stimulation of PnC-projecting CamKIIα-ChR2-mCherry^+^ PPTg fibers evokes excitatory responses in PnC giant neurons in vitro. **(A)** Sample traces showing evoked EPSCs recorded (whole cell patch clamp) in a PnC giant neuron in response to the blue-light photo-stimulation of CamKIIα-ChR2-mCherry^+^ PPTg fibers in PnC sections. **(B)** Sample traces showing evoked EPSPs recorded in a PnC giant neuron in response to the blue-light photo-stimulation of CamKIIα-ChR2-mCherry^+^ PPTg fibers (top traces), in response to paired photo-stimulations separated by 50 ms (middle traces) or 100 ms (bottom traces). Red traces represent the sum of black traces. Representative of N = 10 mice; n = 10 slices.

Aside from giant neurons, the PnC also contains GlyT2^+^ PnC inhibitory neurons previously shown to contribute to PPI (Cano et al., 2021; Huang et al., 2024a). Therefore next, we tested if PPTg glutamatergic neurons also innervate GlyT2^+^ PnC neurons. To do so, we used WT mice injected with the AAVDJ-CamKIIα-mCherry viral vector in the PPTg to transduce PPTg glutamatergic cells with eYFP and a fluorescent parvalbumin (PV^+^) antibody in PnC sections to label PnC inhibitory neurons. Interestingly, eYFP^+^ PPTg glutamatergic fibers were found within the PnC in close apposition to PV^+^ neurons (**Fig. 8**), suggesting that PPTg neurons may be able to activate inhibitory neurons within the PnC to induce PPI at certain ISIs.

**Figure 8.**
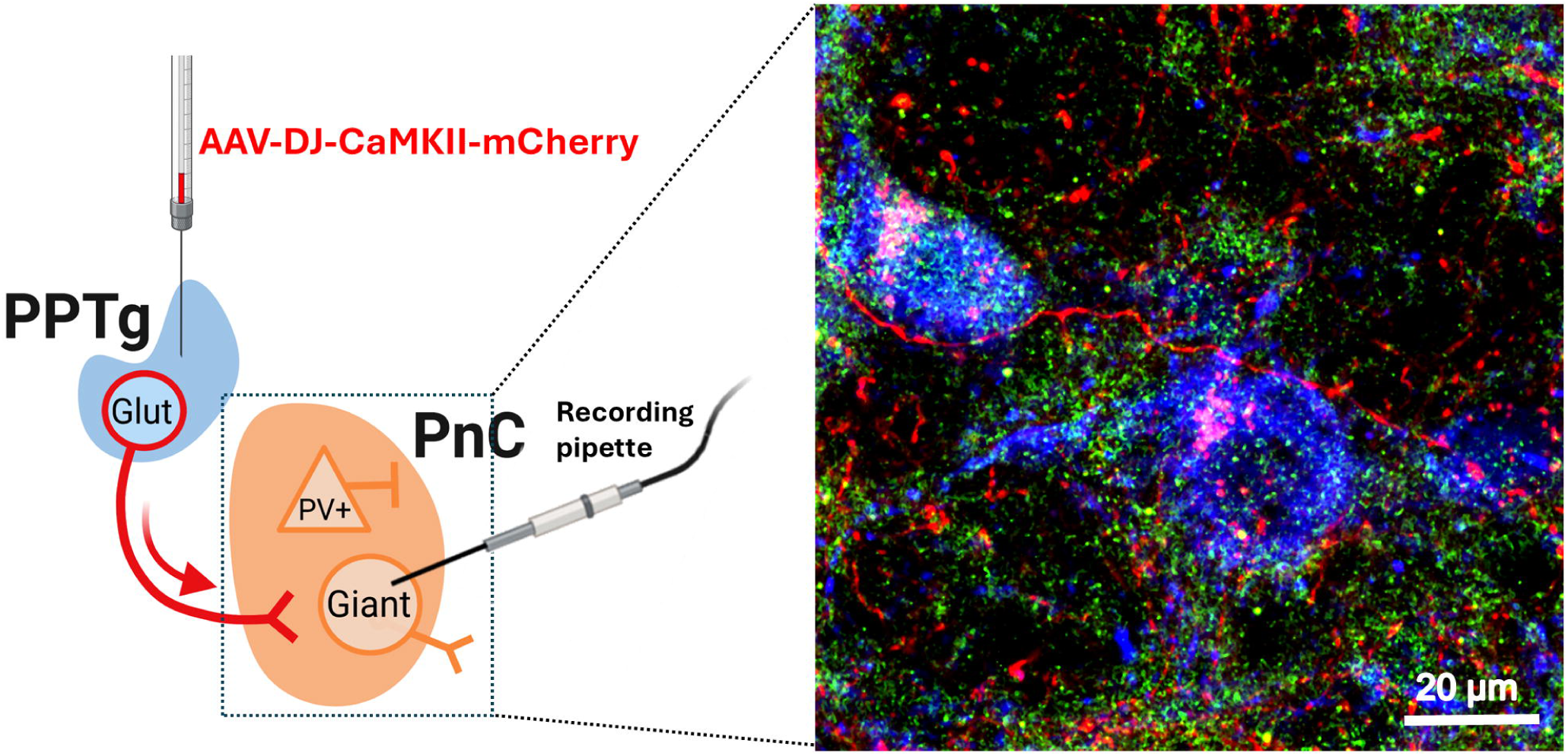
PnC-projecting PPTg fibers course within parvalbumin-postive (PV^+^) PnC neurons. In a WT mouse unilaterally injected with AAV-DJ-CaMKIIα-mCherry in the PPTg, mCherry^+^ PPTg fibers (red) course within PnC PV^+^ neurons (blue) and and are closely apposed to excitatory synapses (PSD95; green), suggesting direct innervation. Image shown at 60x magnification. Image is representative of N = 3 mice.

## Discussion

PPI deficits are linked to cognitive symptoms, but the cellular mechanisms remain ill defined. While the PPTg is clearly involved, we specifically investigated the role of PPTg glutamatergic neurons. Tract tracing and immunohistochemical analyses confirmed that these neurons project to the PnC. Optogenetic inhibition of PPTg-PnC glutamatergic synapses increased acoustic PPI. While optogenetic activation did not affect PPI, activating PPTg-PnC synapses <100 ms before a startling stimulus reduced startle, whereas activation >100 ms prior enhanced startle. *In vitro* optogenetic and electrophysiological single cell recordings revealed that PPTg glutamatergic inputs can activate PnC giant neurons but interestingly, *post hoc* staining also shows that these glutamatergic inputs course in proximity to PnC inhibitory neurons. These findings support a feed-forward and time-dependent excitatory mechanism by which PPTg glutamatergic inputs can bi-directionally modulate the PnC startle circuit. Understanding how the PPTg contributes to PPI is crucial to addressing sensorimotor gating disruptions in neuropsychiatric disorders and improving cognitive treatments.

### Neuroanatomical properties of the PPTg and its glutamatergic neurons

Research has increasingly focused on the PPTg due to its pivotal role in cognitive and sensorimotor functions impaired in diseases like Parkinson’s disease, schizophrenia, and Tourette Syndrome. As a result, the PPTg has emerged as a potential target for deep brain stimulation, showing promise in treating the cognitive symptoms of neurological and psychiatric disorders. Its extensive connections with basal ganglia nuclei enable the relay of activity to thalamic, brainstem, and spinal effectors, integral to motor behavior regulation, including PPI. In fact, while output fibers ascending to cortical and subcortical regions, include the basal ganglia, thalamus, hypothalamus, VTA, amygdala, and frontal cortex (Ryczko and Dubuc, 2013; Roseberry et al., 2016; Lavoie and Parent, 1994c; Dautan et al., 2014; Saper and Loewy, 1982; Edley and Graybiel, 1983; Jackson and Crossman, 1983; Rye et al., 1987; Woolf and Butcher, 1989; Lavoie and Parent, 1994a), descending axons target brainstem nuclei, influencing spinal motor control. Recent studies highlight the role of descending PPTg glutamatergic neurons (Ryczko et al., 2016), particularly those expressing Vglut2, in locomotion and muscle tone regulation. These neurons project ipsilaterally to motor-related brainstem nuclei, forming excitatory synapses in the pontine reticular nucleus (Liang et al., 2012; Caggiano et al., 2018; Huang et al., 2024b). Similarly here, we show that PPTg glutamatergic fibers project mostly unilaterally to the PnC where PSD95 appositions confirm excitatory synapses.

The PPTg contains diverse neuronal populations, including cholinergic, glutamatergic, and GABAergic neurons (Lavoie and Parent, 1994b; Martinez-Gonzalez et al., 2012), with some cholinergic neurons also co-releasing glutamate in rats (Wang and Morales, 2009). Further anatomical studies are needed to precisely define PPTg postsynaptic targets in the PnC. This research could improve understanding of how specific PPTg subpopulations and their postsynaptic targets are altered in diseases associated with PPI deficits.

### Electrophysiological properties of PPTg neurons

The distinct firing patterns of PPTg cholinergic, GABAergic, and glutamatergic neurons across physiological states remain poorly understood. So far, *post hoc* immunohistochemical analyses and intracellular electrophysiological studies in rats have grouped PPTg neurons in three different categories: Type I neurons are small-to-medium fusiform or triangular cells with 3–5 primary dendrites and likely glutamatergic. They exhibit T-type calcium channels-mediated low-threshold spikes and bursting discharges in response to depolarization. Type II neurons, 50% which were ChAT^+^, are medium-to-large polygonal cells with 5–7 primary dendrites. They display A-type potassium currents (*I_A_*) that delay action potentials following hyperpolarization and a tonic, non-bursting pattern. Type III neurons are ChAT^-^ and display mixed characteristics exhibiting both T-type and *I_A_* currents (Kang and Kitai, 1990; Leonard and Llinas, 1990; Takakusaki et al., 1996; Takakusaki and Kitai, 1997; Takakusaki et al., 1997). From our *in vitro* electrophysiology data, CamKIIα^+^ glutamatergic neurons exhibit a spontaneous bursting firing pattern like the type I neurons described above, and efficiently followed a 20 Hz blue-light stimulation train, similar to vGluT2^+^ neurons recorded in vGluT2^Cre^ mice (Huang et al., 2024b).

Interestingly, the intrinsic properties we recorded were similar between CamKIIα^+^ and CamKIIα^-^ PPTg neurons. This discrepancy might arise because CamKIIα^-^ PPTg cells include neurons that co-release acetylcholine (ACh) and glutamate. In fact, ultrastructural studies suggest that ACh and glutamate can be colocalized in axon terminals originating from the mesopontine tegmentum (Clarke et al., 1996; Clarke et al., 1997). Additionally, the action potential half width and the tonic firing pattern of our CamKIIα^-^ PPTg neurons are similar to PPTg cholinergic neurons recently described in mice (Chen and Evans, 2024). Further experiments are needed to characterize the unique electrophysiological properties of mouse CamKIIα^-^PPTg neurons that are purely cholinergic, GABAergic or ChAT/vGlut2^+^.

Our *in vitro* electrophysiology findings also suggest that PPTg glutamatergic neurons likely project outside the PPTg (Kroeger et al., 2017) and target regions such as the striatum (Assous et al., 2019) and PnC (Caggiano et al., 2018) to modulate locomotion. Striatum-projecting PPTg glutamatergic neurons have been studied in detail, but less is known about the PnC-projecting ones. Optogenetic activation of PPTg glutamatergic axons induces monosynaptic excitatory responses in fast-spiking GABAergic interneurons driving feedforward inhibition in the striatum. Therefore, based on our neuroanatomical data, it is tempting to suggest that PPTg glutamatergic neurons also drives feedforward inhibition that can contribute to PPI by activating PnC PV^+^ inhibitory neurons. Such mechanism could be parallel to PPTg glutamatergic fibers monosynaptically activating PnC giant neurons, to provide an excitation/inhibition balance within the PnC startle circuit. The PnC houses glutamatergic giant neurons and PV^+^ neurons that include GlyT2^+^ glycinergic and GABAergic neurons. We previously showed that GlyT2^+^ PnC neurons contribute to PPI and can be activated by CeA glutamatergic inputs (Cano et al., 2021; Huang et al., 2024a). Since inputs to the PnC also arise from the PPTg, future electrophysiological studies should clarify how PPTg glutamatergic neurons target different postsynaptic elements within the PnC to affect PPI.

### The contribution of PPTg glutamatergic neurons to locomotion and startle modulation

Previous mouse studies have demonstrated that vGluT2^+^ PPTg neurons can enhance (Caggiano et al., 2018; Masini and Kiehn, 2022), inhibit (Josset et al., 2018; Dautan et al., 2021), or exert a combined effect on locomotion (Carvalho et al., 2020). The variability in these outcomes likely arises from the activation of distinct pathways involving PPTg glutamatergic neurons, each with specific downstream targets. For example, activating a subset of MLR vGluT2^+^ neurons that project to the substantia nigra halts locomotion, while activating projections to the spinal cord induces body extension in mice and those projecting to the medulla enhance locomotion (Ferreira-Pinto et al., 2021; Goñi-Erro et al., 2023). Similarly, Huang et al. (2024b) reported that activating all vGluT2^+^ PPTg neurons or only those projecting to the PnC inhibits locomotion, whereas activation of vGluT2^+^ PPTg neurons projecting to the zona incerta facilitates movement. These findings provide strong evidence for the presence of functionally distinct PPTg glutamatergic neuron populations with unique postsynaptic targets.

Ample evidence also supports the contribution of the PPTg to PPI in rodents. Early studies suggested that PPTg cholinergic neurons play a key role in mediating PPI since cholinergic-specific lesions in the rodent PPTg disrupted PPI (Koch et al., 1993; Bosch and Schmid, 2008). However, recent research challenged this view as optogenetic activation and chemogenetic inhibition of these neurons in rodents did not alter acoustic PPI responses but rather PPTg cholinergic neurons were shown to facilitate startle responses (MacLaren et al., 2014; Azzopardi et al., 2018; Fulcher et al., 2020). Additionally, PPTg cholinergic boutons observed in close proximity to both glutamatergic and GABAergic cells in the inferior colliculus, suggest that ACh may modulate both excitatory and inhibitory circuits (Noftz et al., 2020). However, the postsynaptic targets of PPTg cholinergic neurons in the PnC remain unclear, and the mechanism by which these cholinergic neurons enhance startle is still unknown.

Since few studies focused on the involvement of non-cholinergic neuronal populations in PPI modulation, we focused on the contribution of PPTg glutamatergic neurons and demonstrated that they project to PnC giant excitatory neurons and PV^+^ inhibitory neurons. Our *in vivo* photo-inhibition experiments confirm that PPTg glutamatergic neurons provide an excitatory drive to the PnC during PPI. Furthermore, our photo-activation experiments revealed additional insights into how PPTg glutamatergic neurons can modulate the PnC startle circuit. Like cholinergic neurons, activation of PPTg glutamatergic neurons enhanced startle responses at longer interstimulus intervals, likely by activating PnC giant neurons. In contrast, activation of these PPTg glutamatergic neurons at shorter interstimulus intervals reduced startle (mimicking PPI), possibly through the activation of PV^+^ inhibitory neurons. Future experiments should determine whether the PPTg contains functionally distinct glutamatergic neuron subpopulations that target different neuronal circuits during PPI.

## Acknowledgements

We thank Ms. Andrea Gouin for the technical support and for the animal care; Dr. Joseph A. Gogos (Columbia University), Dr. Amy B. MacDermott (Columbia University), Dr. Mary Torregrossa (University of Pittsburgh), Dr. David Moorman (UMass Amherst) and Dr. Joseph Bergan (UMass Amherst) for the insightful discussions; and Dr. James Chambers for the assistance with the three-dimensional reconstructions performed at the Light Microscopy Facility at UMass Amherst. Figures were created with BioRender.com

---

**Supplementary Figure 1. Retrograde labeling of PnC-projecting PPTg neurons.**

**(A)** Schematic illustrating the unilateral injection of AAVrg-CAG-tdTomato in the PnC of WT mice. **(B)** Representative PnC microinjection site of AAVrg-CAG-tdTomato shown at 10x magnification. **(C)** Retrogradely labeled PPTg neuronal cell bodies express td-Tomato (red) and a subset of them co-express Choline acetyltransferase (ChAT; green) or GAD67 (cyan), sown at 10x magnification. **(D)** Close up view of labeled PPTg neurons from the boxed area shown in (C). Images are representative of N = 3 animals. Scale bars: B, C = 1mm; D = 100μm.

**Supplementary Figure 2. CamKIIα-eYFP expression and ChAT immuno-reactivity in the PPTg.**

**(A, B)** Representative PPTg coronal section showing CamKIIα-eYFP expression (green) and immunofluorescence of the cholinergic marker ChAT (red), which delineates the PPTg. **(C)** Overlay of A and B. **(D, E)** eYFP^+^ and ChAT^+^ PPTg cell bodies shown at higher magnification. **(F)** Overlay of D and E showing that a few PPTg cells co-express eYFP and ChAT (white arrow). Representative of N = 4 mice.

**Supplementary figure 3. Intrinsic and synaptic properties of CamKIIα-ChR2-mCherry^+^ and CamKIIα-ChR2-mCherry^-^ PPTg neurons.**

**(A)** Schematic illustrating the in vitro whole cell patch clamp recording experiments performed in ChR2-mCherry^+^ (red) and ChR2-mCherry^-^ (non-fluorescent, black) PPTg neurons of WT mice. Also shown are representative spontaneous excitatory post synaptic current (sEPSC) traces obtained from whole cell voltage clamp recordings of ChR2-mCherry^+^ (blue trace) or non-fluorescent (black trace) PPTg neurons. **(B)** Plot showing the cumulative distribution of sEPSC amplitude of ChR2-mCherry^+^ (blue) or non-fluorescent (black) PPTg neurons. **Inset**, spontaneous firing pattern of ChR2-mCherry^+^ (blue) or non-fluorescent (black) PPTg neurons. **(C)** Plot of the firing rate of ChR2-mCherry^+^ (open circles) or non-fluorescent (black circles) PPTg neurons as a function of depolarizing currents. **Inset**, representative voltage traces. **(D)** Representative blue light-evoked EPSPs (top) and EPSCs (bottom) recorded in the absence (black traces) or presence of glutamate receptor blockers CNQX and AP5 (red traces). **(E)** Graph showing the mean paired-pulse ratio of blue light-evoked EPSPs as a function of interpulse intervals (IPIs) ranging from 10 to 1000 ms, in the absence (black circles) or presence of glutamate receptor blockers CNQX and AP5 (red circles). **(F)** Graph showing the mean paired-pulse ratio of blue light-evoked EPSCs as a function of IPIs ranging from 10 to 1000ms, in the absence (black circles) or presence of glutamate receptor blockers CNQX and AP5 (red circles). N=10 mice; n = 10 slices. Data represented as mean ± SEM. *P>0.05

## Notes

### Competing Interest Statement

The authors have declared no competing interest.

