## Supplemental Information for "Midbrain Glutamatergic Neurons Modulate the Acoustic Startle Reflex and Prepulse Inhibition in Mice"

**Abbreviated title:** Midbrain Glutamatergic Neurons Modulate ASR and PPI

**Author Names and Affiliations:** Luis Enrique Martinetti^2,4^, Erika Correll^1,4^, Adolfo Ernesto Cuadra^1^, Gina Castellano^3^, and Karine Fénelon^1^*

^1^: Biology Department, University of Massachusetts Amherst, Life Science Laboratories, 240 Thatcher Road, Amherst, MA, 01002, U.S.A.

^2^: Department of Biological Sciences, University of Texas at El Paso, 500 West University Avenue, El Paso, TX, 79912, U.S.A.

^3^: Commonwealth Honors College, University of Massachusetts Amherst, 157 Commonwealth Avenue, Amherst, MA, 01002, U.S.A.

^4^: These authors contributed equally

**Supplemental Information**

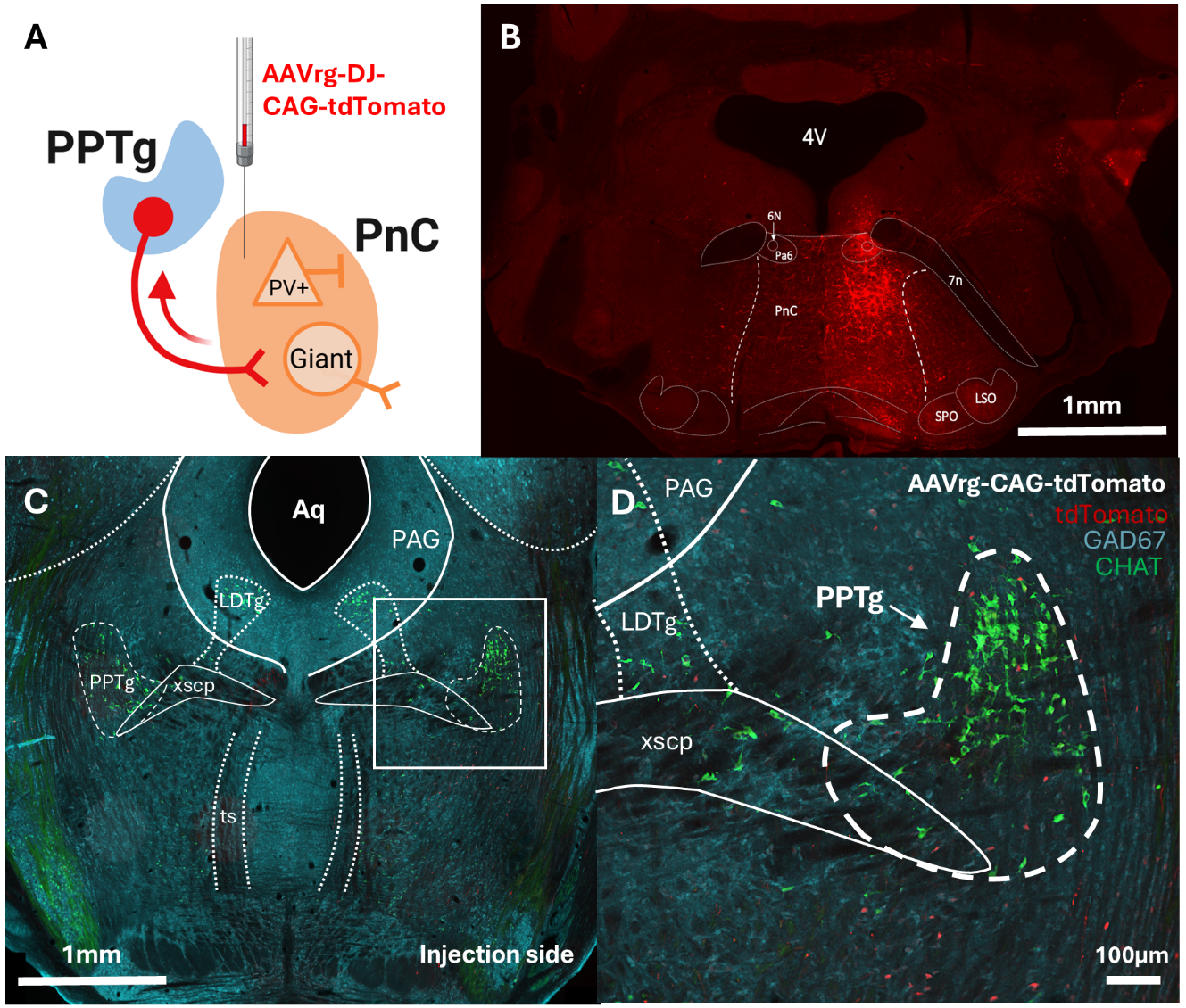

**Supplementary Figure 1. Retrograde labeling of PnC-projecting PPTg neurons.**

**(A)** Schematic illustrating the unilateral injection of AAVrg-CAG-tdTomato in the PnC of WT mice. **(B)** Representative PnC microinjection site of AAVrg-CAG-tdTomato shown at 10x magnification. **(C)** Retrogradely labeled PPTg neuronal cell bodies express td-Tomato (red) and a subset of them co-express Choline acetyltransferase (ChAT; green) or GAD67 (cyan), sown at 10x magnification. **(D)** Close up view of labeled PPTg neurons from the boxed area shown in (C). Images are representative of N = 3 animals. Scale bars: B, C = 1mm; D = 100μm

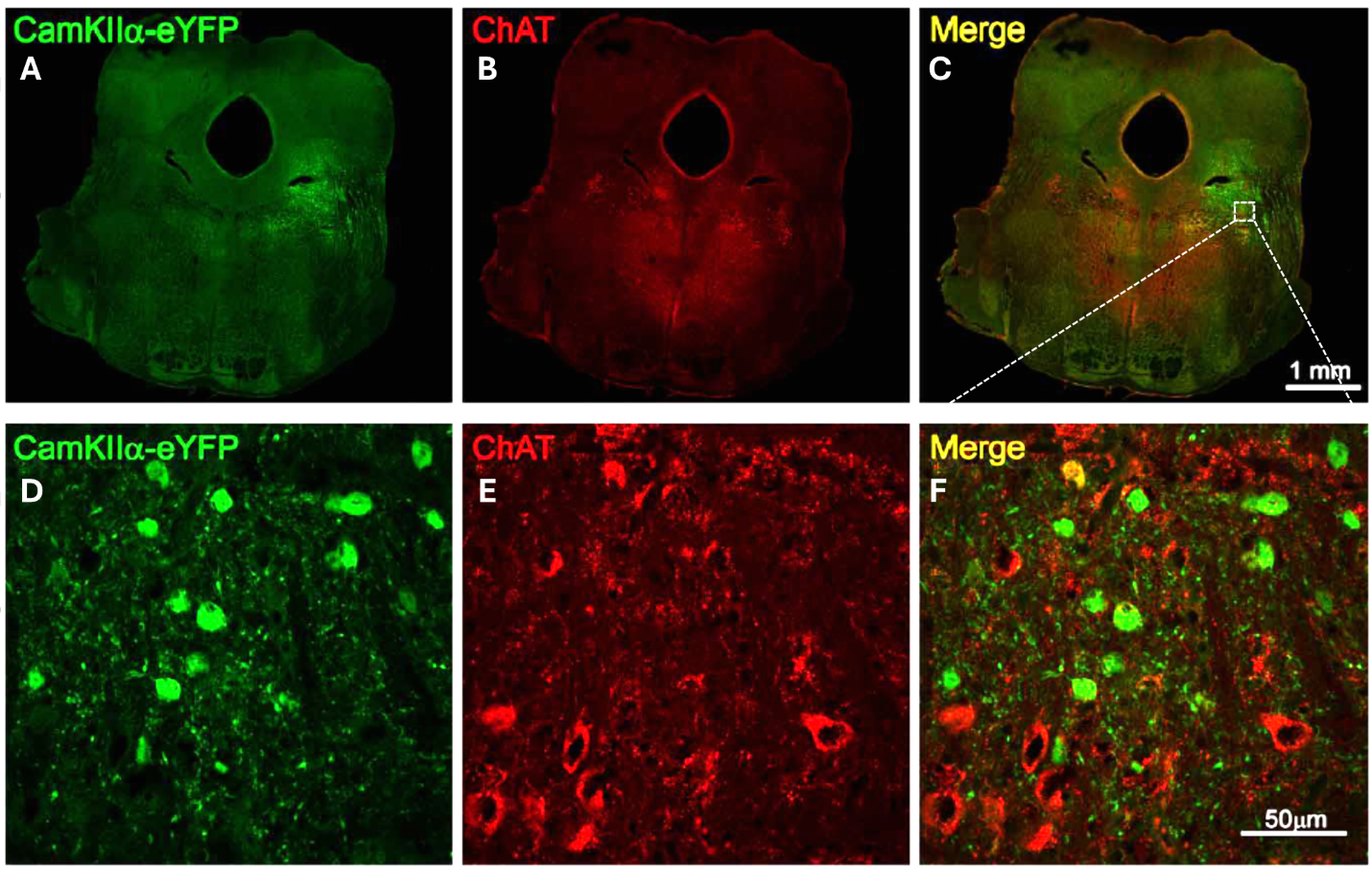

**Supplementary Figure 2. CamKIIα-eYFP expression and ChAT immuno-reactivity in the PPTg.**

**(A, B)** Representative PPTg coronal section showing CamKIIα-eYFP expression (green) and immunofluorescence of the cholinergic marker ChAT (red), which delineates the PPTg. **(C)** Overlay of A and B. **(D, E)** eYFP^+^ and ChAT^+^ PPTg cell bodies shown at higher magnification. **(F)** Overlay of D and E showing that a few PPTg cells co-express eYFP and ChAT (white arrow). Representative of N = 4 mice.

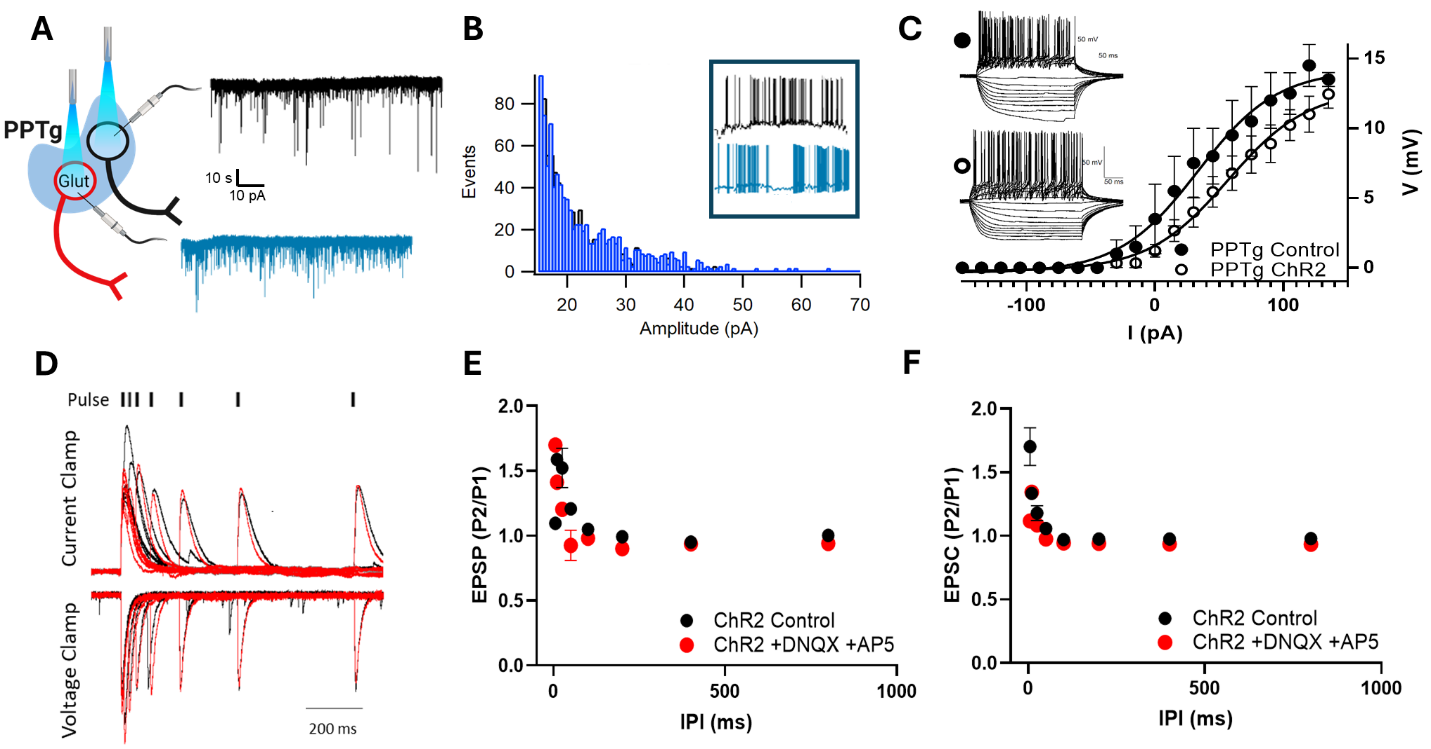

**Supplementary figure 3. Intrinsic and synaptic properties of CamKIIα-ChR2-mCherry^+^ and CamKIIα-ChR2-mCherry^-^ PPTg neurons.**

**(A)** Schematic illustrating the in vitro whole cell patch clamp recording experiments performed in ChR2-mCherry^+^ (red) and ChR2-mCherry^-^ (non-fluorescent, black) PPTg neurons of WT mice. Also shown are representative spontaneous excitatory post synaptic current (sEPSC) traces obtained from whole cell voltage clamp recordings of ChR2-mCherry^+^ (blue trace) or non-fluorescent (black trace) PPTg neurons. **(B)** Plot showing the cumulative distribution of sEPSC amplitude of ChR2-mCherry^+^ (blue) or non-fluorescent (black) PPTg neurons. **Inset**, spontaneous firing pattern of ChR2-mCherry^+^ (blue) or non-fluorescent (black) PPTg neurons. **(C)** Plot of the firing rate of ChR2-mCherry^+^ (open circles) or non-fluorescent (black circles) PPTg neurons as a function of depolarizing currents. **Inset**, representative voltage traces. **(D)** Representative blue light-evoked EPSPs (top) and EPSCs (bottom) recorded in the absence (black traces) or presence of glutamate receptor blockers CNQX and AP5 (red traces). **(E)** Graph showing the mean paired-pulse ratio of blue light-evoked EPSPs as a function of interpulse intervals (IPIs) ranging from 10 to 1000 ms, in the absence (black circles) or presence of glutamate receptor blockers CNQX and AP5 (red circles). **(F)** Graph showing the mean paired-pulse ratio of blue light-evoked EPSCs as a function of IPIs ranging from 10 to 1000ms, in the absence (black circles) or presence of glutamate receptor blockers CNQX and AP5 (red circles). N=10 mice; n = 10 slices. Data represented as mean ± SEM. *P>0.05
